# SemVac: A Semantic Vaccinology Paradigm Powered by LLMs for Antigen Discovery

**DOI:** 10.64898/2026.07.13.737696

**Authors:** Yunxiang Zhao, You Shu, Linfeng Shu, Peng Lv, Xiangyang Chi, Dongsheng Li, Jun Zhang, Zhen Huang, Hongguang Ren, Junjie Xu, Xiaodong Zai, Wei Chen

## Abstract

Reverse vaccinology identifies vaccine antigens from pathogen genomes, yet existing methods rely mainly on sequence and structure and overlook the functional and immunological knowledge recorded in the published literature. We introduce semantic vaccinology, a paradigm in which large language models (LLMs) reason over literature-derived protein descriptions to predict protective antigens. Implemented as SemVac, the workflow retrieves publications linked to each protein through PaperBLAST, condenses the evidence into a structured semantic profile, and prompts an LLM to return an antigenicity probability. Benchmarked against a curated 246-protein bacterial benchmark and the specialized protein-language and geometric-deep-learning predictor PLGDL, the best of 14 general-purpose LLMs matched or exceeded the precision of PLGDL; the open-weight Kimi K2 0905 offered the strongest performance-cost balance. Predictions were robust to masking of vaccine keywords, reproducible across repeated inference, and generalized to a 1,200-protein cross-pathogen dataset. Surprisingly, explicit chain-of-thought reasoning increased recall but lowered precision in every model tested, revealing over-reasoning in biological scoring. Applied to the mpox virus proteome, SemVac recovered the established mpox antigen repertoire and prioritized uncharacterized candidates. Two of these, A30L and C19L, received independent experimental support in recent orthopoxvirus vaccine development, providing external validation. For one candidate, B20R, the model generated a coherent but false TNF-decoy narrative unsupported by curated annotations, demonstrating that LLM confabulation can be detected when reasoning traces are cross-checked against curated resources. Semantic vaccinology therefore establishes the literature as an explicit, auditable third modality alongside sequence and structure, while making its failure modes transparent and correctable.

## INTRODUCTION

Vaccines remain the most effective tool for preventing infectious disease, but their development is limited by the difficulty of choosing the right antigen from thousands of proteins in a pathogen genome [1,2]. Reverse vaccinology addressed this problem by reading genomes directly: starting from the whole genome of *serogroup B Neisseria meningitidis*, proteins were ranked by sequence features such as surface exposure, secretion and adhesin probability [1,3]. Computational support for reverse vaccinology has passed through three generations. Early tools filtered whole proteomes with hard thresholds on sequence-predicted properties—subcellular localization, signal peptides, transmembrane topology and adhesin-likeness [4–6]; machine-learning methods then replaced those thresholds with trained classifiers while keeping the features largely unchanged, combining the same biological annotations with evolutionary conservation and physicochemical descriptors [7–9]. Both generations rest on hand-crafted features and therefore inherit the error of the annotation tools that produce them [10]. The current generation removes that bottleneck in two ways. Protein language models trained at evolutionary scale supply embeddings in which structure and function emerge without supervision [11,12], and because antibody accessibility is governed by the folded protein rather than the linear sequence, geometric deep-learning architectures now learn directly from predicted three-dimensional coordinates, which high-accuracy structure prediction has made available for essentially any protein [13,14]. Recent frameworks combine both channels—sequence embeddings with a graph neural network over predicted structures—supporting residue-level epitope interpretation and proteome-wide candidate prioritization [15], and in our own work leading to the experimental identification of a new protective mpox antigen, G10R [16].

Yet a protein’s value as a vaccine target is not fully captured by its sequence or fold. Whether it mediates adhesion, invasion, virulence or immune evasion, and whether it is accessible to antibodies during infection, are properties described predominantly in the biomedical literature [17,18]. Curated databases such as Protegen [17] and the Immune Epitope Database [18] capture parts of this knowledge, but the functional narrative for each protein is distributed across tens of thousands of publications and integrated mainly through expert effort. No computational framework has automated this evidence layer.

Large language models now encode substantial biomedical knowledge, support literature-grounded reasoning in medicine and public health, and are beginning to reshape how scientists interact with biological data [19–22]. Benchmarks such as LAB-Bench measure their capacity for biological research [23], and autonomous research agents have embedded language models in complete discovery pipelines [24]. In vaccinology, an antigen-centric knowledge graph that fuses large-scale literature mining with LLM-augmented reasoning has improved antigen selection for tuberculosis vaccines, suggesting that structured literature evidence can guide computational antigen discovery [25].

Whether general-purpose LLMs can match specialized predictors by reasoning directly over unstructured protein descriptions has, however, not been tested. We formalize this idea as semantic vaccinology: the prediction of vaccine candidacy from literature-derived semantic representations of proteins through LLM reasoning. We position the paradigm as a third modality alongside sequence-centric and structure-aware reverse vaccinology, and we implement it as SemVac (Fig. 1). We benchmark 14 general-purpose LLMs against a curated bacterial antigen set and the specialist predictor PLGDL [16], analyze the effect of explicit reasoning on scoring behavior, and apply the workflow to the complete proteome of mpox virus, a WHO-priority pathogen for which additional vaccine antigens are actively sought [26,27]. The study is a computational, proof-of-concept analysis; no new experiments were performed, and all candidate rankings remain hypotheses pending laboratory validation.

**Fig. 1.**
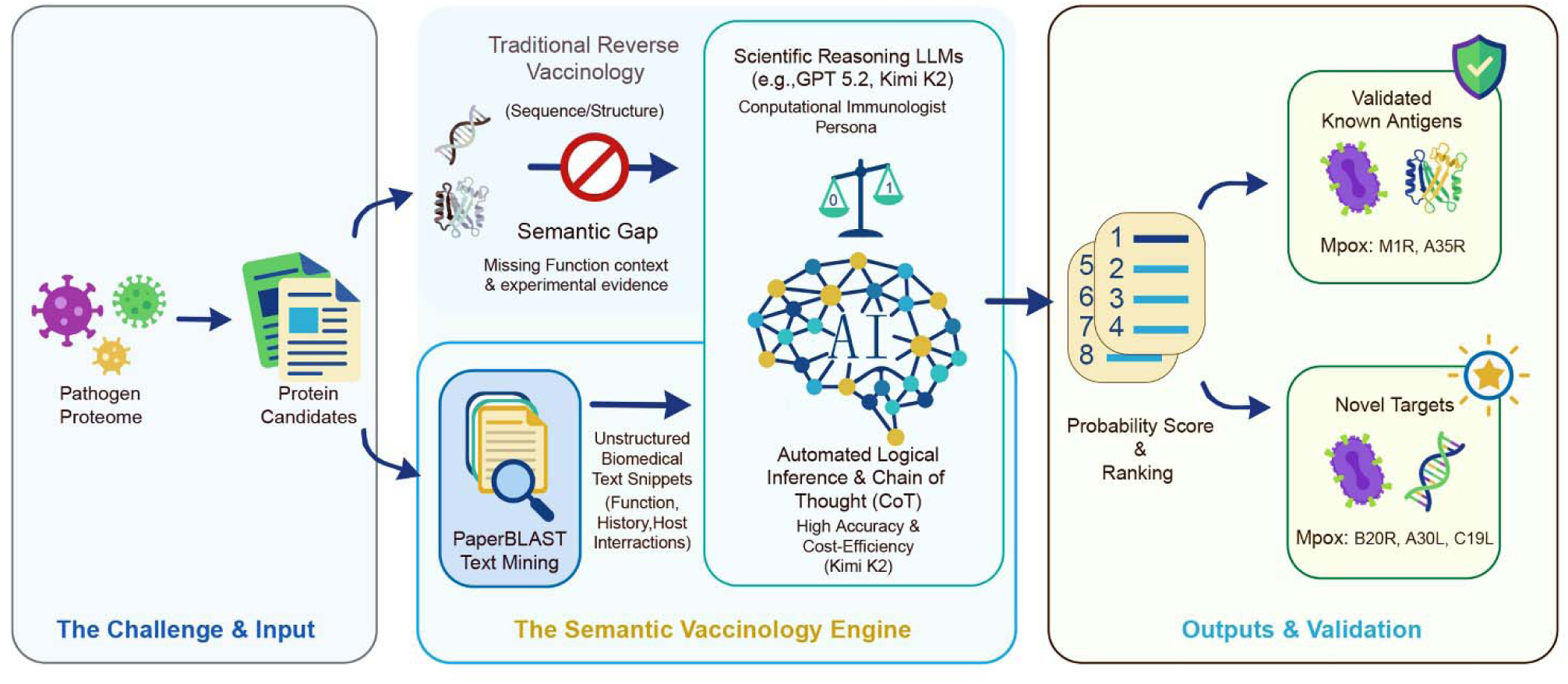
The SemVac workflow for literature-derived antigen prediction. Protein sequences from a pathogen proteome are queried against PaperBLAST to retrieve functionally relevant literature snippets, which are organized into structured semantic evidence profiles and evaluated by a large language model to yield an antigenicity probability. The workflow makes the evidence layer explicit and auditable.

## RESULTS

### SemVac computes literature-grounded antigenicity scores

Semantic vaccinology replaces engineered sequence features with literature-derived semantic evidence as the input to antigen prediction (Fig. 1). Its premise is that most of what is known about a pathogen protein already exists in prose rather than in any database field. Realizing that premise is difficult. The knowledge is not missing but unanchored: it is scattered across a biomedical literature of tens of millions of articles, in which a single protein may be denoted by a gene name, a locus tag, several database accessions and informal synonyms, and in which naming conventions differ among pathogens, strains and research communities. PaperBLAST [28] supplies the bridge, and SemVac is built on it. PaperBLAST searches the full text of articles in Europe PMC for protein and gene identifiers and links each mention to a protein sequence, and it additionally incorporates curated resources (Swiss-Prot, GeneRIF and EcoCyc) in which sequences and publications are already paired. The result is a precomputed correspondence between sequences and the literature, which we enter through homology.

For each protein in a pathogen proteome, SemVac queries PaperBLAST [28] by protein-protein BLAST and aggregates the publications linked to the query and its homologues (expect value < 0.001). On average 268 publications were retrieved per protein (median 122) (Supplementary Fig. S2). Text segments describing function, localization, virulence association, secretion, immune relevance and experimental observations were extracted and organized into a structured semantic evidence profile (Supplementary Fig. S1). Each profile was presented to an LLM instructed to act as an expert computational immunologist and to return a single probability between 0 and 1 that the protein is a protective vaccine antigen.

### Leading general-purpose LLMs match or exceed the specialized predictor PLGDL

To test whether an LLM can convert literature-derived semantics into accurate antigen judgements, we evaluated SemVac on the independent bacterial protective antigen benchmark (iBPA) [9], with its composition detailed in Materials and Methods (Supplementary Fig. S3). Labels rest on immunization and challenge experiments in animal models rather than on computational annotation, which makes the set stringent for comparing reverse vaccinology methods. It also raises an immediate concern: the experiments that define the labels are part of the same literature SemVac reads. We address this directly below.

Closed-source frontier models defined the upper precision band (Fig. 2A). GPT-5.2 Pro achieved the highest precision at the 0.5 threshold (0.897), followed by GPT-5.2 (0.883) and Claude Opus 4.5 (0.854). The open-weight model Kimi K2 0905 ranked fourth overall (0.838), ahead of every other open model and ahead of PLGDL [16] (0.799), a purpose-built predictor that integrates evolutionary-scale sequence modeling with geometric reasoning. Qwen 3 Max (0.788), GLM-5 (0.761), Grok 4.1 Fast (0.756) and DeepSeek V3.2 (0.750) occupied intermediate positions. Positive and negative score distributions were sharply separated for the leading models, as quantified by the area under the receiver operating characteristic curve (ROC-AUC) of 0.915 for GPT-5.2 Pro, 0.914 for Kimi K2 0905 and 0.940 for Claude Opus 4.5, the highest among all evaluated systems (Fig. 2B; Supplementary Fig. S5). The separation was likewise evident in violin comparisons (Fig. 2C; P = 2.85E-29 and P = 6.27E-30; Supplementary Fig. S6) and in confusion matrices with balanced error structures at the 0.5 threshold (Fig. 2D; Supplementary Fig. S7). Full metrics are provided in Supplementary Table S1.

**Fig. 2.**
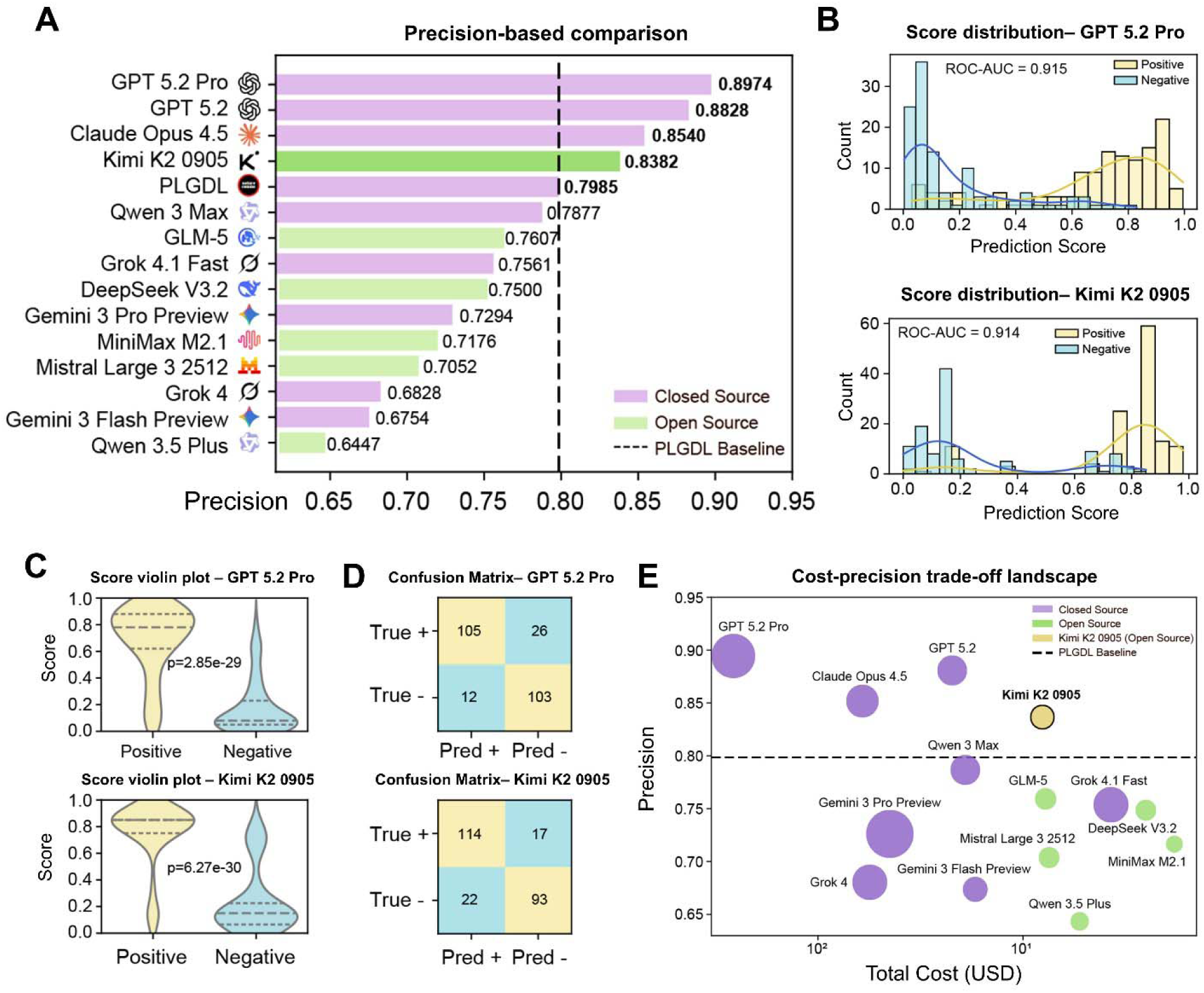
Benchmark performance and cost-precision landscape of SemVac across 14 LLMs on the 246-protein bacterial antigen set. (A) Precision at the 0.5 threshold; the dashed line marks the non-LLM specialist predictor PLGDL (0.7985). Purple, closed-source models; green, open-weight models. (B) Predicted antigenicity score distributions with kernel density estimates for positive (yellow) and negative (blue) samples from GPT-5.2 Pro (ROC-AUC = 0.915) and Kimi K2 0905 (ROC-AUC = 0.914). (C) Antigenicity-score violin plots for positive versus negative proteins (GPT-5.2 Pro: P = 2.85E-29; Kimi K2 0905: P = 6.27E-30). (D) Confusion matrices at the 0.5 threshold for GPT-5.2 Pro and Kimi K2 0905. (E) Cost-precision trade-off landscape (total inference cost in US dollars, log scale; calculated from official API pricing applied to total token consumption, including reasoning tokens); the dashed line marks PLGDL.

Practical deployment depends on inference cost as much as on accuracy. We computed per-model expenditure from official API pricing applied to total token consumption, including reasoning tokens. Closed-source models were substantially more expensive: GPT-5.2 Pro consumed US$258.77 for the 246-protein benchmark, 32- to 142-fold more than open-weight alternatives (US$1.82-8.04; e.g., MiniMax M2.1, DeepSeek V3.2 and Kimi K2 0905). The cost-precision landscape separated the two classes cleanly, with closed models occupying the high-cost, high-precision region and open models achieving competitive precision at a small fraction of the expense (Fig. 2E). Kimi K2 0905 offered the most favorable balance, retaining fourth place in precision at roughly 3% of the cost of the top-ranked model.

### The semantic signal is reproducible, model-independent and pathogen-agnostic

Predictions were highly correlated within model families, yet the ranking did not split simply by vendor (Fig. 3A,B). The pairwise Pearson correlation matrix grouped closely related models together, and the score heatmap showed broad agreement on which proteins score high and which score low; open- and closed-weight systems nonetheless clustered together rather than separating by access type. Two conclusions follow. First, architecturally distinct models converge on comparable evaluations of the same semantic evidence, so the performance reported above is not an artifact of one tokenizer, training corpus or alignment procedure. Second, because these models were trained independently and are exposed only to the same literature profiles, the shared component of their predictions must reside in the evidence itself rather than in a model-specific cue. None of the tested models was trained for antigen prediction; their agreement therefore reflects the transfer of literature-embedded biological knowledge to a specialized inference task.

**Fig. 3.**
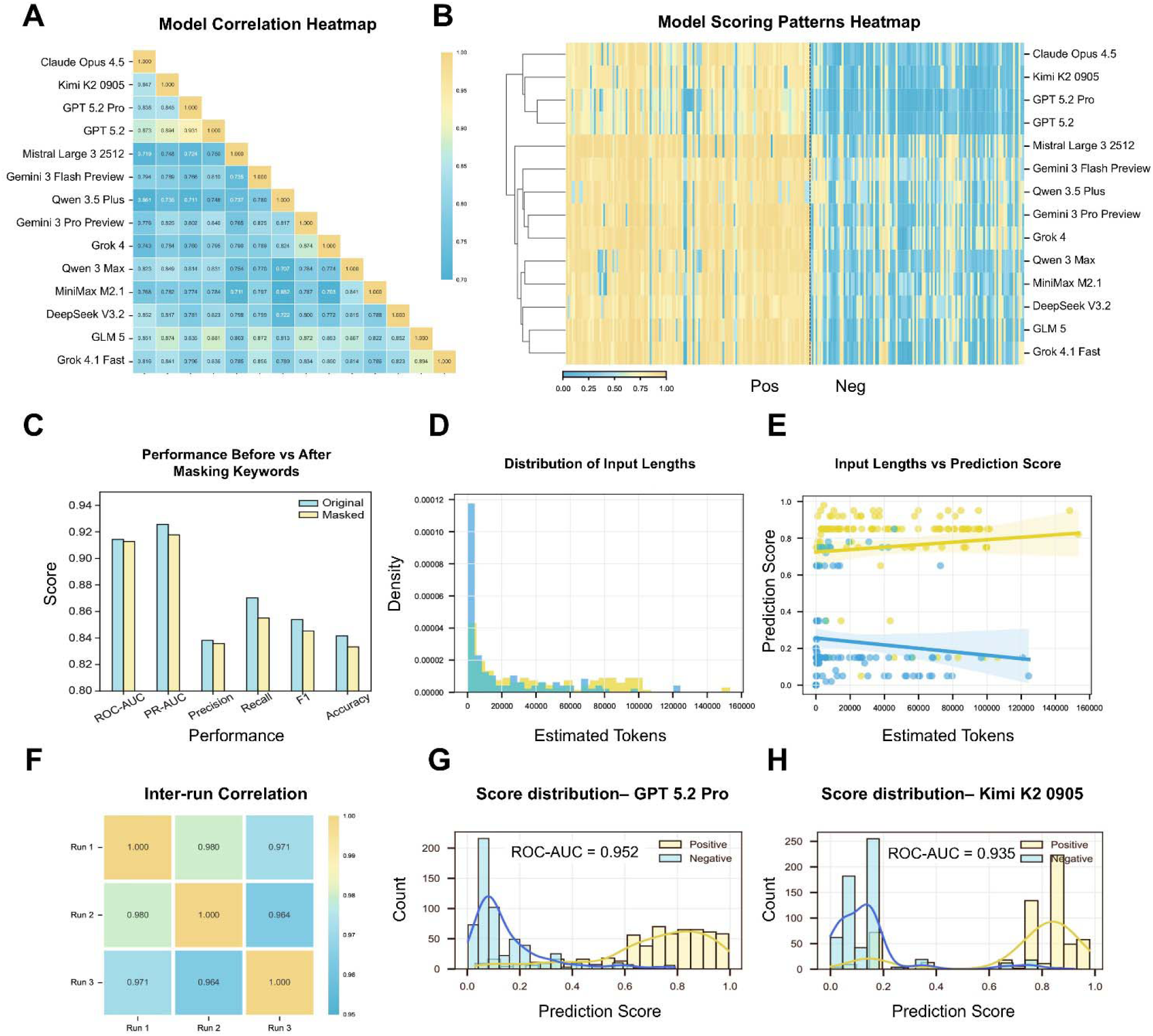
Robustness, reproducibility and cross-pathogen generalization. (A) Score heatmap of all evaluated LLMs. The horizontal axis represents proteins and the vertical axis represents models. (B) Pairwise Pearson correlation heatmap of antigenicity scores across LLMs on the benchmark dataset. (C) Performance of Kimi K2 0905 before (Original) and after (Masked) removing every publication containing vaccine-related keywords. (D) Distribution of estimated input lengths in tokens. (E) Input length versus predicted antigenicity score for Kimi K2 0905 (positive: Pearson r = 0.103; negative: r = -0.104). (F) Inter-run correlation matrix across three independent repeats of the complete benchmark (pairwise correlations of key metrics between 0.971 and 1.000). (G-H) Antigenicity score histograms for GPT-5.2 Pro (G) and Kimi K2 0905 (H) on the 1,200-protein cross-pathogen dataset.

A central concern for literature-driven prediction is that models may simply match explicit mentions of vaccines, protection or antigens in the retrieved text. We tested this directly by removing every publication containing such keywords from the evidence profiles before inference with Kimi K2 0905. Precision declined only marginally (0.838 before versus 0.836 after masking; Fig. 3C), indicating that the model’s judgements are anchored in the substance of protein characterization literature rather than in superficial keyword matching.

Repeated inference produced highly consistent scores across three independent runs of the complete benchmark, with pairwise correlations of precision, F1 and area under the precision-recall curve (PR-AUC) between 0.971 and 1.000 (Fig. 3F), and input length was only weakly associated with predicted scores (Fig. 3D,E), together supporting the stability of the evaluation framework.

To assess whether SemVac generalizes beyond bacterial antigens, we assembled an extended dataset of 600 positive and 600 negative proteins spanning viral, bacterial and eukaryotic antigens from Protegen [17], IEDB [18] and published reverse vaccinology screens [16] (Supplementary Fig. S4). Both GPT-5.2 Pro and Kimi K2 0905 maintained clearly separated score distributions across this cross-pathogen set, with precision of 0.938 and 0.933 and ROC-AUC of 0.952 and 0.935, respectively (Fig. 3G,H and Supplementary Table S2). Semantic reasoning therefore transfers across taxonomic groups and antigen classes without pathogen-specific adjustment.

### An over-reasoning effect in antigen scoring

Many frontier LLMs expose an explicit reasoning mode that generates step-by-step chain-of-thought before answering [29,30]. Because judging an antigen looks like a task that should benefit from careful step-by-step thinking, we asked whether such modes improve it. Across all four models that support both configurations, reasoning modes produced a consistent pattern: recall increased in every case (by 8.40, 6.11, 1.53 and 0.77 percentage points for Claude Opus 4.5, Kimi K2 0905, DeepSeek V3.2 and Gemini 3 Flash Preview, respectively), whereas precision decreased in every case (by 0.63, 7.57, 2.75 and 1.21 percentage points for Claude Opus 4.5, Kimi K2 0905, DeepSeek V3.2 and Gemini 3 Flash Preview, respectively) (Fig. 4A,B and Supplementary Table S3). Discrimination measured by ROC-AUC improved slightly, indicating that reasoning mode inflates scores globally rather than sharpening them: models become more inclusive, not more selective.

**Fig. 4.**
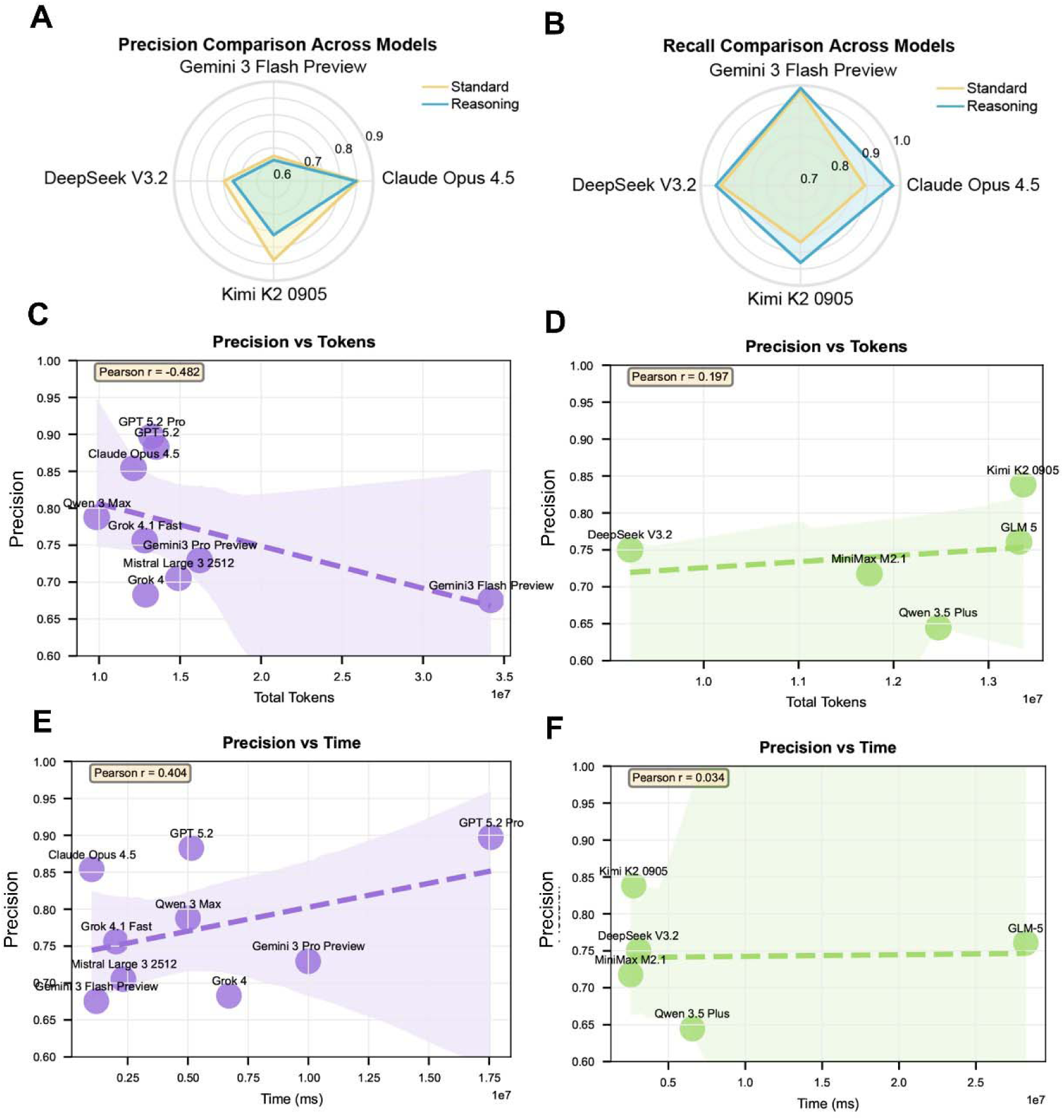
Time, token trade-offs and the effect of explicit reasoning. (A-B) Radar plots of precision (A) and recall (B) under standard versus reasoning-enabled modes for the four models supporting both configurations. Reasoning mode consistently increased recall and decreased precision. (C-D) Precision versus total token consumption for closed-source (C; Pearson r = -0.482) and open-weight (D; r = 0.197) models. (E-F) Precision versus wall-clock inference time for closed-source (E; r = 0.404) and open-weight (F; r = 0.034) models.

Inspection of reasoning traces suggests a mechanism. Verbose step-by-step deduction encourages the model to elaborate biologically plausible but unfounded narratives, such as hallucinated mechanisms or overinterpreted homology, which convert borderline evidence into confident positive scores. The effect is not a simple consequence of longer inputs, because input length was only weakly associated with predicted scores (Fig. 3E). Our observations parallel recent reports that extended deliberation can degrade performance on tasks that reward concise pattern recognition [31–33] and that reasoning-heavy biomedical questions can be more brittle than knowledge-heavy ones [34]. Biological discovery problems are governed by evolutionary and physicochemical constraints that manifest as recurring patterns rather than lengthy deductive chains; forcing extended verbal reasoning appears to amplify the generation of plausible-sounding but unsupported explanations. Consistent with this interpretation, Kimi K2 0905, an architecture that does not rely on explicit long-form reasoning, achieved the best open-weight performance without requiring an explicit reasoning mode. We do not read these results as an argument against reasoning in biology, but as evidence that the form of reasoning matters: representation-level inference acquired during pretraining may align better with the structure of biological knowledge than explicit verbal deduction, at least for scoring tasks of this kind.

Analyses of wall-clock time and token usage revealed no significant associations with performance: precision showed only a weak, non-significant trend with time consumption among closed models (Pearson r = 0.404; Fig. 4E) and none among open models (r = 0.034; Fig. 4F), whereas token usage showed weak negative and positive trends in the two groups, respectively (closed models r = -0.482; open models r = 0.197; Fig. 4C,D). None of these correlations reached significance given the small number of models, but together they indicate diminishing returns on inference expenditure. Open-weight LLMs thus constitute practical, scalable instruments for semantic vaccinology, particularly in academic and resource-limited settings.

### Proteome-wide application to mpox virus

Mpox virus remains a public health priority after the 2022 outbreaks and continued spread of clade I viruses in Africa [27]. Although modified vaccinia Ankara and, more recently, mRNA vaccines encoding the M1R, A35R, B6R, A29L and H3L antigens provide protection [35–37], the antigenic space of the virus has been explored only partially [38,39], and orthogonal discovery methods continue to reveal new protective targets [39]. We therefore applied SemVac, with Kimi K2 0905 as the inference engine, to the complete reference proteome of mpox virus strain MPXV_USA_2022_MA001 (ON563414.3, clade IIb). Predicted antigenicity scores spanned a wide dynamic range (0.05-0.92), with a low-scoring majority and a distinct high-scoring tail (Fig. 5A).

**Fig. 5.**
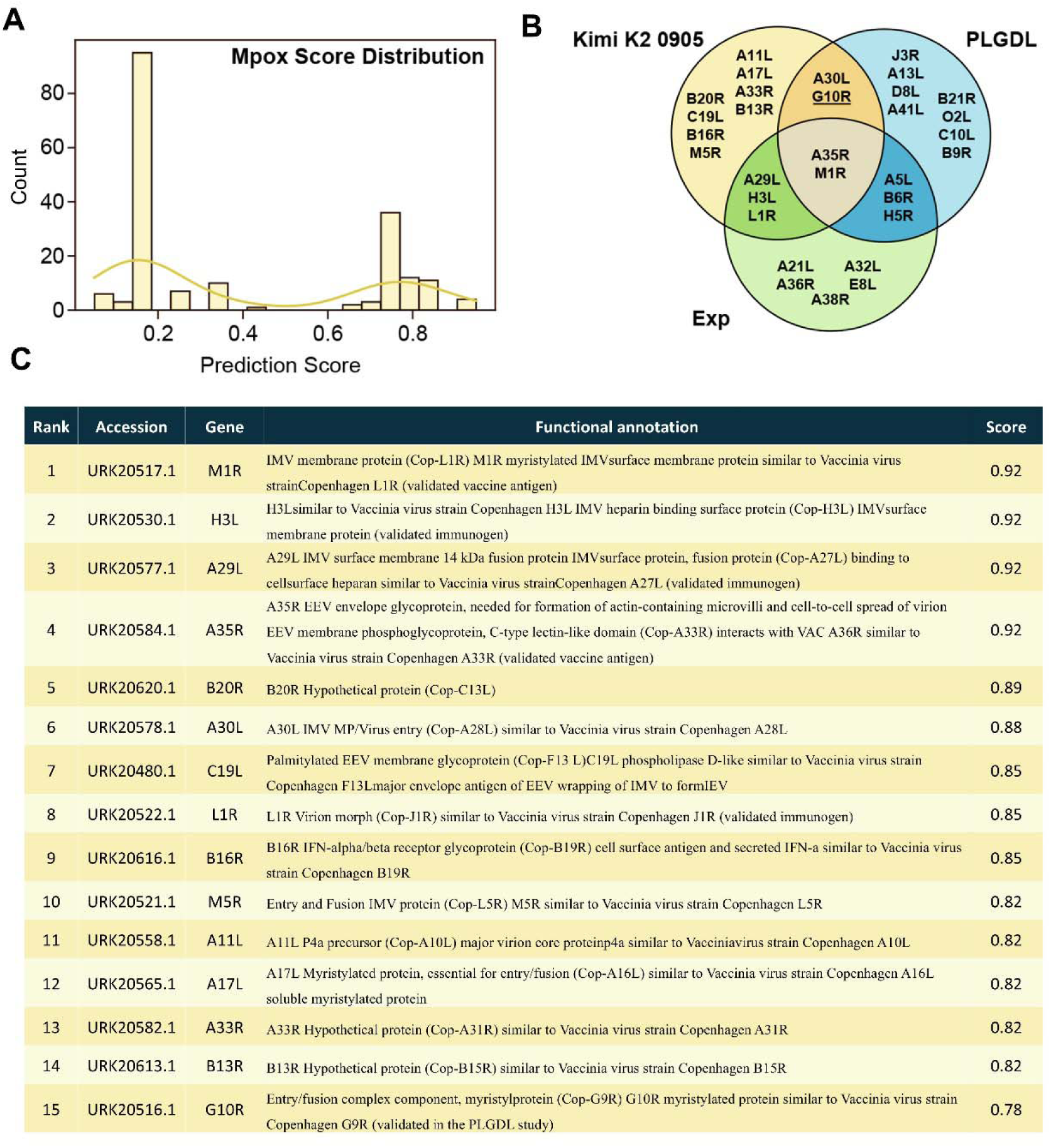
Proteome-wide application of SemVac to mpox virus. (A) Antigenicity score distribution across the 190 proteins of the MPXV_USA_2022_MA001 proteome predicted by Kimi K2 0905 (range 0.05-0.92). (B) Overlap between the top 15 SemVac candidates, structure-based PLGDL predictions and experimentally validated antigens (Exp). (C) Top 15 SemVac-ranked candidates with accessions, genes and functional annotations. The asterisks mark the two candidates (A30L and C19L) that received independent experimental support in recent orthopoxvirus vaccine development [40–42].

SemVac recovered the established antigen repertoire with high fidelity (Fig. 5B). Several experimentally validated vaccine antigens, M1R, A35R, A29L, H3L and L1R, ranked within the top 15, and four candidates, namely M1R, A35R, A30L and G10R, overlapped with the predictions of the structure-based PLGDL framework [16], among which G10R had been experimentally confirmed as a protective antigen in the PLGDL study itself. This convergence of independent modalities, one textual and one structural, constitutes a form of cross-validation for both methods. The remaining top candidates (B20R, C19L, B16R, M5R, A11L, A17L, A33R and B13R) had not been prioritized by sequence- or structure-based approaches and represent newly prioritized targets (Fig. 5C).

Strikingly, recent orthopoxvirus vaccine-development efforts provided experimental support for two top-ranked candidates. A30L, whose vaccinia orthologue A28L is a component of the virion entry-fusion complex, was also prioritized among the candidate antigens evaluated in systematic profiling of mpox antigens for next-generation mRNA vaccine development [40, 41]. C19L, the mpox homologue of the vaccinia major envelope protein F13L (p37) of the extracellular enveloped virion (EEV), was likewise evaluated among the extracellular-virion antigens in a subsequent systematic screen [41]; moreover, a monovalent C19L-encoding mRNA vaccine has been reported to confer complete protection against lethal ectromelia-virus challenge in mice, supporting its use as a pan-orthopoxvirus vaccine antigen [42]. The patent data were generated well before the construction of the semantic literature database underpinning SemVac and reside outside the academic literature corpus it indexes, and both of the supporting studies were published after its temporal cutoff (December 2025); the convergence therefore constitutes a de facto external validation. This independent convergence strongly suggests that the literature signal captured by SemVac corresponds to biologically meaningful antigenic features. Full candidate annotations and orthologue assignments are given in Supplementary Table S4.

### An inspectable failure mode: B20R functional confabulation

Inspection of the model’s reasoning traces illustrates how semantic evidence is synthesized and where it can fail (Fig. 6A). A30L (97.95% similar to its vaccinia orthologue A28L) was prioritized on the basis of its essential role in membrane fusion, consistent with curated annotation as an entry-fusion-complex component [43]. C19L (99.46% similar to vaccinia F13L, p37) was prioritized through its essential role in virion morphogenesis and extracellular enveloped virus formation, consistent with its status as the target of the antiviral drug tecovirimat [38,44]. The third trace, for the novel candidate B20R (96.84% similar to its vaccinia orthologue C13L), documents a limit of the paradigm as plainly as its strengths. The model assembled genuine evidence—strong orthopoxvirus conservation, predicted structural exposure and a high score (0.89)—yet its functional narrative attributed secreted tumor-necrosis-factor-decoy activity to B20R, an annotation that curated resources do not support: the established mpox TNF decoy receptor is the CrmB protein encoded by J2L [45,46].

**Fig. 6.**
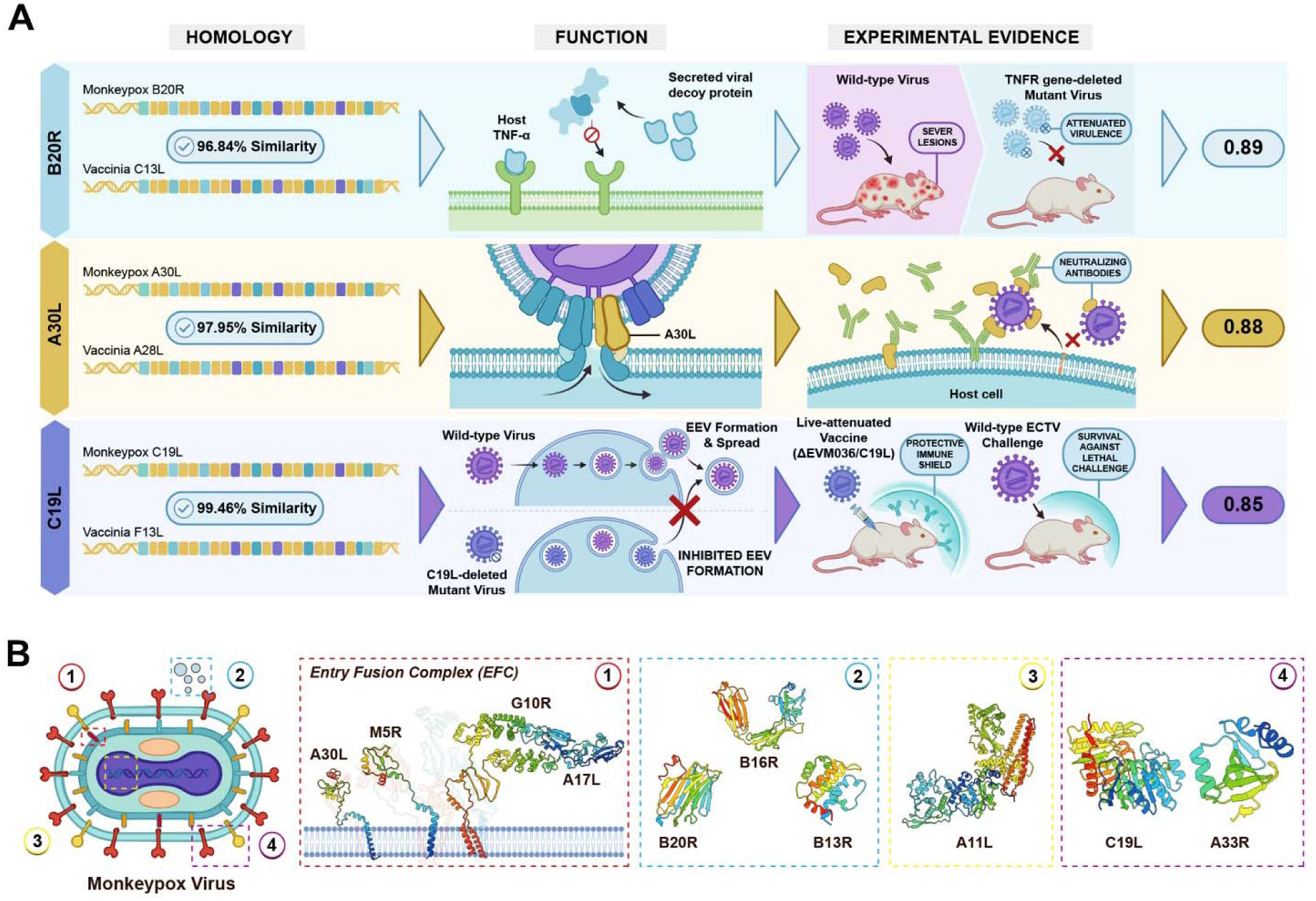
Model reasoning traces and structural context of top mpox candidates. (A) Reasoning traces for B20R, A30L and C19L, each integrating sequence homology to the vaccinia orthologue (similarity shown), functional annotation and experimental literature evidence, with the final antigenicity score at right (B20R 0.89, A30L 0.88, C19L 0.85). The function panel for B20R documents a functional misattribution: the model attributes secreted TNF-decoy activity that curated annotation does not support (the established mpox TNF decoy is CrmB, encoded by J2L), whereas the A30L and C19L narratives agree with curated annotation. Model-generated rationales are hypotheses pending experimental verification. (B) mpox virion architecture and the entry-fusion complex, showing predicted localization of candidate antigens; predicted structures are shown for novel candidates.

This is not a hypothetical risk but a concrete, inspectable instance of functional misattribution, in which homology-centered reasoning completed a missing annotation with a wrong yet biologically coherent story. We therefore treat every model-generated functional narrative as a hypothesis and cross-check all candidate annotations against curated orthologue assignments [43]. Genomic and structural localization of the remaining unvalidated candidates placed most of them within or adjacent to the virion membrane and entry-fusion machinery (Fig. 6B), consistent with physical accessibility to antibodies.

## DISCUSSION

This work establishes the scientific literature as a third modality for computational antigen discovery. Sequence-based reverse vaccinology reads the genome; structure-aware methods read the fold; semantic vaccinology reads the published record of what proteins do. That the leading general-purpose LLMs, without any task-specific training, matched or exceeded a purpose-built predictor in precision indicates that the knowledge required for antigen evaluation is already embedded in frontier models and can be elicited through structured evidence presentation. The concurrent demonstration that an antigen-centric knowledge graph, fusing large-scale literature mining with LLM-augmented reasoning, supports antigen selection for tuberculosis vaccines [25] points to the same principle from an independent direction, suggesting that literature-grounded inference is a generalizable foundation for vaccinology rather than a peculiarity of one implementation. Our quantitative, task-grounded benchmark also complements expert-centered assessments of artificial-intelligence cognitive abilities in systems vaccinology [47]: whereas those evaluations probe what models can reason about, SemVac measures, with reproducible scoring and cost accounting, how reliably general-purpose models execute one concrete discovery step. This semantic channel complements the sequence- and structure-based channels that already drive machine-learning antigen and epitope prediction: the same targets that literature reasoning prioritizes can subsequently be scored by these predictors, so that the three modalities together span discovery, prioritization and engineering [16,48,49].

The over-reasoning pitfall deserves emphasis because it runs counter to the prevailing assumption that more reasoning is better. In every model tested, explicit chain-of-thought made predictions more inclusive and less precise. This mirrors the overthinking phenomenon documented for large reasoning models on simple problems [31] and the broader observation that test-time computation must be matched to task structure [32,33]. For evidence-synthesis tasks of the kind studied here, extended verbal deduction appears to manufacture confidence rather than accuracy. Practically, we recommend standard inference modes for antigen scoring, reserving explicit reasoning for hypothesis narration in which its generative elaboration is an asset rather than a liability.

Semantic predictions inherit the noise of the literature and of database annotation, and our own application surfaced a concrete instance rather than an abstract risk. Asked about B20R, the model produced a confident, internally coherent narrative that misattributed TNF-decoy functionality to a protein whose curated annotation carries no immune-evasion role (Fig. 6A). The failure mode is characteristic of LLM reasoning: where evidence is sparse, homology-centered inference tends to complete the gap with the most biologically plausible story, and the fluency of the output makes the error hard to detect from the narrative alone. Three observations temper the concern. First, the error was detectable within seconds by cross-checking curated orthologue assignments [43], a step costing little compared with experimental screening and one we now regard as mandatory for any literature-driven prediction pipeline. Second, the misattribution did not corrupt the score itself, which was driven by conservation and exposure evidence rather than by the faulty annotation. Third, the workflow’s explicit evidence layer makes such failures inspectable: the reasoning trace preserves exactly which assertions the model made and on what grounds. Scores and narratives must therefore be read separately—the former as a prioritization signal, the latter as a hypothesis.

Several limitations qualify our findings. The benchmark is bacterial and modest in size; labels derive from experimental immunization and challenge data compiled from the literature and may embed publication bias; absence of protective evidence is not proof of non-protection; proteins with sparse literature coverage are disadvantaged by construction; and model behaviors will drift as vendors update their systems, although the robustness of the framework across 14 models and two datasets mitigates version-specific concerns. The cross-pathogen set, while taxonomically broader, reuses the same literature-consensus label source and was not independently re-validated experimentally. Prospective experimental testing of candidates such as A30L and C19L is required before any claim of protective efficacy; the B20R misattribution (Fig. 6A) illustrates why model narratives must remain hypotheses until then.

Looking forward, the static one-shot inference used here is the simplest instantiation of semantic vaccinology. The field is moving toward closed discovery loops in which candidate predictions are experimentally tested and the results fed back for refinement, exemplified by multi-agent frameworks that generate, critique and rank hypotheses for experimental validation [50,51]. Embedding semantic vaccinology in such agent-based frameworks would close the loop between literature-grounded prediction and experimental feedback. Integration with sequence- and structure-based predictors as complementary evidence channels, proteome-wide application to viral and eukaryotic pathogens, extension to cancer neoantigen discovery, and systematic prospective testing of candidates are natural next steps. More broadly, the same principle applies wherever expert judgement depends on synthesizing unstructured literature, including drug target identification, protein function annotation and host-pathogen interaction analysis.

## MATERIALS AND METHODS

### Benchmark datasets

The benchmark was derived from the independent bacterial protective antigen benchmark (iBPA) assembled by Ong et al. [9], comprises 131 protective antigens and 115 non-protective proteins (246 in total) curated from published reverse vaccinology resources. Protective antigens were defined as proteins with experimental validation of protective efficacy in animal immunization or challenge models; non-protective samples were bacterial proteins lacking evidence of protective immunity despite experimental evaluation. An extended validation set of 1,200 proteins (600 positive, 600 negative) spanning viral, bacterial and eukaryotic antigens was assembled from Protegen [17], IEDB [18] and published reverse vaccinology screens [16]; duplicate and highly redundant records were removed before evaluation. All sequences were standardized to FASTA format and associated with UniProt identifiers where available. Supplementary Fig. S3 shows the taxonomic distribution of the bacterial benchmark; Supplementary Fig. S4 shows that of the extended dataset.

### Literature retrieval and semantic evidence construction

Structured semantic evidence profiles were generated for each protein using PaperBLAST [28]. For every query sequence, homologous proteins with linked publications were identified through protein-protein BLAST against the PaperBLAST database (NCBI BLAST+; database snapshot as of December 2025), retaining matches with an expect value below 0.001. All linked publications were aggregated (mean 268, median 122 publications per protein; Supplementary Fig. S2), and text segments describing protein function, localization, virulence association, secretion characteristics, immune relevance, pathogenicity and experimental observations were extracted automatically. The resulting corpus was organized into a profile containing protein identifiers and annotation metadata, homology statistics and alignment confidence, source organism information, literature-derived functional descriptions and experimentally reported immunological evidence.

### Prompt formulation and model inference

A unified prompt framework was used for all models, consisting of a system-level instruction and a protein-specific user input. The system instruction defined the model’s role as an expert computational immunologist and vaccinologist and required integration of the retrieved literature evidence with general immunological principles. User inputs were formatted as structured objects containing protein identifiers, homology statistics, publication metadata and extracted snippets. All LLMs were accessed through the OpenAI-compatible interface of OpenRouter, each required to output a score between 0.0 and 1.0 representing the predicted probability that the protein is a protective antigen, with the score provided at the end of the model’s response. Decoding temperature was set to 0. All evaluations were conducted between January and March 2026. Supplementary Fig. S1 shows a representative prompt. The evaluated models are listed in Supplementary Table S1.

### Reasoning-mode configuration

For models supporting explicit reasoning or chain-of-thought inference, reasoning mode was activated through the corresponding OpenRouter thinking parameters, allowing the model to allocate additional reasoning tokens before producing the final output [30]. All other settings, including prompts, input formatting, temperature and output constraints, were identical to the standard protocol, and models were still required to return only a single probability value.

### Evaluation metrics and cost accounting

Predicted probabilities were compared with ground-truth labels to compute standard binary classification metrics: precision (TP/(TP+FP)), recall (TP/(TP+FN)), F1 score (harmonic mean of precision and recall) and accuracy ((TP+TN)/(TP+TN+FP+FN)). A score threshold of 0.5 was used for confusion matrices and threshold-dependent metrics. ROC-AUC and PR-AUC were computed across all thresholds. Precision was designated the primary metric given the cost of experimentally validating false-positive vaccine candidates. Inference cost was calculated from official API pricing applied to total token consumption, including reasoning tokens, and is reported in US dollars. Wall-clock time and prompt, completion and reasoning token counts were recorded for every model. For reproducibility, the complete benchmark was scored three times independently with Kimi K2 0905, and pairwise Pearson correlations of precision, F1 score and PR-AUC were computed across runs. The influence of input length on prediction behavior was assessed by correlating the estimated token length of each literature input with the predicted score, separately for positive and negative samples.

### mpox proteome analysis

The complete reference proteome of mpox virus strain MPXV_USA_2022_MA001 was obtained from NCBI (ON563414.3) and processed with the identical SemVac pipeline used for the benchmark. Candidate annotations and orthologue assignments were cross-checked against curated mpox virus–vaccinia virus orthologue mappings and UniProt [52] and InterPro [53] family annotations; functional claims for novel candidates are reported as hypotheses accordingly.

## ACKNOWLEDGMENTS

We thank colleagues at the National Key Laboratory of Advanced Biotechnology and the College of Computer, National University of Defense Technology, for discussion. This study was supported by the National Natural Science Foundation of China (32571110, 62421002 and 62576357).

During the preparation of this manuscript the authors used GLM-5.3 (Zhipu AI) for text polishing and error checking; the scientific conclusions, data analysis and experimental design were made by the authors.

## Author contributions

W.C., X.Z., J.X. and H.R. conceived and supervised the project. Y.Z., Y.S. and L.S., as co-first authors, performed the data analysis and processing, including benchmark evaluation, robustness and reproducibility analyses, cost accounting, proteome-wide mpox screening and interpretation of the results. P.L. and X.C. performed the biological and orthologue-annotation analyses. D.L., J.Z. and Z.H. contributed to data curation. X.Z., Y.Z., Y.S. and L.S. wrote the manuscript with input from all authors. All authors approved the final version.

## Competing interests

The authors declare no competing interests.

## DATA AVAILABILITY

The benchmark datasets and the results required to reproduce the figures of this study are available on Zenodo at https://zenodo.org/records/19384187. Source code of the SemVac workflow is available at the same record.

## Notes

### Competing Interest Statement

The authors have declared no competing interest.

### Summary of Updates

This version of the manuscript has been substantially revised in structure, scope and interpretation, while the benchmark datasets, evaluated models and primary quantitative results remain unchanged. Structure and figures. The main text has been reorganized from continuous prose into explicit Introduction, Results (with six subsection headings), Discussion and Materials and Methods sections. The original two main figures were expanded and reorganized into six main figures (Fig. 1-6), and the supplementary figures were consolidated (Fig. S1-S7). New robustness analyses. We added (i) a reproducibility assessment based on three independent repeats of the complete benchmark, with pairwise correlations of precision, F1 and PR-AUC between 0.971 and 1.000 (Fig. 3F); (ii) analyses of input length versus predicted scores (Fig. 3D,E); and (iii) wall-clock time and token usage versus performance, indicating diminishing returns on inference expenditure (Fig. 4C-F). External validation of mpox candidates. A new Results section reports that two top-ranked candidates received independent experimental support after our analysis: A30L was prioritized in systematic profiling of mpox antigens for next-generation mRNA vaccine development, and a monovalent C19L mRNA vaccine conferred complete protection against lethal ectromelia virus challenge in mice. Because these data postdate the temporal cutoff of our literature database or reside outside it, this convergence constitutes a de facto external validation (Fig. 5). Reinterpretation of B20R. B20R, previously highlighted as a promising novel candidate linked to immune evasion, is now reported as a concrete, inspectable case of functional confabulation: the model-generated TNF-decoy narrative is not supported by curated annotation, as the established mpox TNF decoy receptor is CrmB encoded by J2L. The Results, Discussion and Fig. 6A were revised accordingly, and cross-checking of all model narratives against curated orthologue assignments is now a mandatory step in the workflow. Updated proteome analysis. The mpox reference proteome was updated from strain Zaire-96-I-16 (187 proteins) to MPXV_USA_2022_MA001 (clade IIb, 190 proteins), candidate annotations were expanded (new Supplementary Table S4), and the bacterial benchmark is now explicitly identified as the iBPA dataset assembled by Ong et al. References and declarations. The reference list was expanded from 28 to 53 entries, an AI-assistance declaration was added, and the author contribution statements were revised.

