## Supplemental Figures and Tables for "SemVac: A Semantic Vaccinology Paradigm Powered by LLMs for Antigen Discovery"

**Supplementary Tables**

**Table S1. Full performance metrics for all 14 evaluated LLMs and the PLGDL baseline on the 246-protein bacterial antigen benchmark, including ROC-AUC, PR-AUC, precision, recall, F1, accuracy and total inference cost.**

| **Rank** | **Model** | **ROC-AUC** | **PR-AUC** | **Precision** | **Recall** | **F1** | **Accuracy** | **Cost (US$)** |
| --- | --- | --- | --- | --- | --- | --- | --- | --- |
| 1 | GPT-5.2 Pro | 0.9150 | 0.9338 | 0.8974 | 0.8015 | 0.8468 | 0.8455 | 258.77 |
| 2 | GPT-5.2 | 0.9335 | 0.9472 | 0.8828 | 0.8626 | 0.8726 | 0.8659 | 22.14 |
| 3 | Claude Opus 4.5 | 0.9401 | 0.9470 | 0.8540 | 0.8931 | 0.8731 | 0.8618 | 60.71 |
| 4 | Kimi K2 0905 | 0.9143 | 0.9256 | 0.8382 | 0.8702 | 0.8539 | 0.8415 | 8.04 |
| 5 | PLGDL | 0.8476 | 0.8561 | 0.7985 | 0.8359 | 0.8168 | 0.8017 | - |
| 6 | Qwen 3 Max | 0.8905 | 0.8959 | 0.7877 | 0.8779 | 0.8303 | 0.8089 | 19.12 |
| 7 | GLM-5 | 0.9357 | 0.9417 | 0.7607 | 0.9466 | 0.8435 | 0.8130 | 7.75 |
| 8 | Grok 4.1 Fast | 0.9174 | 0.9058 | 0.7561 | 0.9466 | 0.8407 | 0.8089 | 3.72 |
| 9 | DeepSeek V3.2 | 0.9113 | 0.9064 | 0.7500 | 0.9389 | 0.8339 | 0.8008 | 2.52 |
| 10 | Gemini 3 Pro Preview | 0.9284 | 0.9347 | 0.7294 | 0.9466 | 0.8239 | 0.7846 | 44.58 |
| 11 | MiniMax M2.1 | 0.9025 | 0.9130 | 0.7176 | 0.9313 | 0.8106 | 0.7683 | 1.82 |
| 12 | Mistral Large 3 (2512) | 0.8987 | 0.9085 | 0.7052 | 0.9313 | 0.8026 | 0.7561 | 7.46 |
| 13 | Grok 4 | 0.9317 | 0.9341 | 0.6828 | 0.9695 | 0.8013 | 0.7439 | 55.89 |
| 14 | Gemini 3 Flash Preview | 0.9093 | 0.8862 | 0.6754 | 0.9847 | 0.8012 | 0.7398 | 17.10 |
| 15 | Qwen 3.5 Plus | 0.8840 | 0.8993 | 0.6447 | 0.9695 | 0.7744 | 0.6992 | 5.29 |

**Table S2. Performance of GPT-5.2 Pro and Kimi K2 0905 on the extended cross-pathogen dataset (600 positive, 600 negative proteins).**

| **Model** | **ROC-AUC** | **PR-AUC** | **Precision** | **Recall** | **F1** | **Accuracy** |
| --- | --- | --- | --- | --- | --- | --- |
| GPT-5.2 Pro | 0.9520 | 0.9561 | 0.9383 | 0.9100 | 0.9239 | 0.9225 |
| Kimi K2 0905 | 0.9347 | 0.9402 | 0.9327 | 0.9050 | 0.9186 | 0.9175 |

**Table S3. Standard versus reasoning-enabled inference modes. Reasoning mode consistently increases recall and reduces precision across all models that support both configurations.**

| **Model** | **Mode** | **ROC-AUC** | **PR-AUC** | **Precision** | **Recall** | **F1** | **Accuracy** |
| --- | --- | --- | --- | --- | --- | --- | --- |
| Claude Opus 4.5 | Standard | 0.9401 | 0.9470 | 0.8540 | 0.8931 | 0.8731 | 0.8618 |
| Claude Opus 4.5 | Reasoning | 0.9566 | 0.9567 | 0.8477 | 0.9771 | 0.9078 | 0.8943 |
| Gemini 3 Flash Preview | Standard | 0.9093 | 0.8862 | 0.6754 | 0.9847 | 0.8012 | 0.7398 |
| Gemini 3 Flash Preview | Reasoning | 0.9384 | 0.9395 | 0.6633 | 0.9924 | 0.7951 | 0.7276 |
| DeepSeek V3.2 | Standard | 0.9113 | 0.9064 | 0.7500 | 0.9389 | 0.8339 | 0.8008 |
| DeepSeek V3.2 | Reasoning | 0.9242 | 0.9266 | 0.7225 | 0.9542 | 0.8224 | 0.7805 |
| Kimi K2 0905 | Standard | 0.9143 | 0.9256 | 0.8382 | 0.8702 | 0.8539 | 0.8415 |
| Kimi K2 0905 | Reasoning | 0.9252 | 0.9372 | 0.7625 | 0.9313 | 0.8385 | 0.8089 |

**Supplementary Figures**

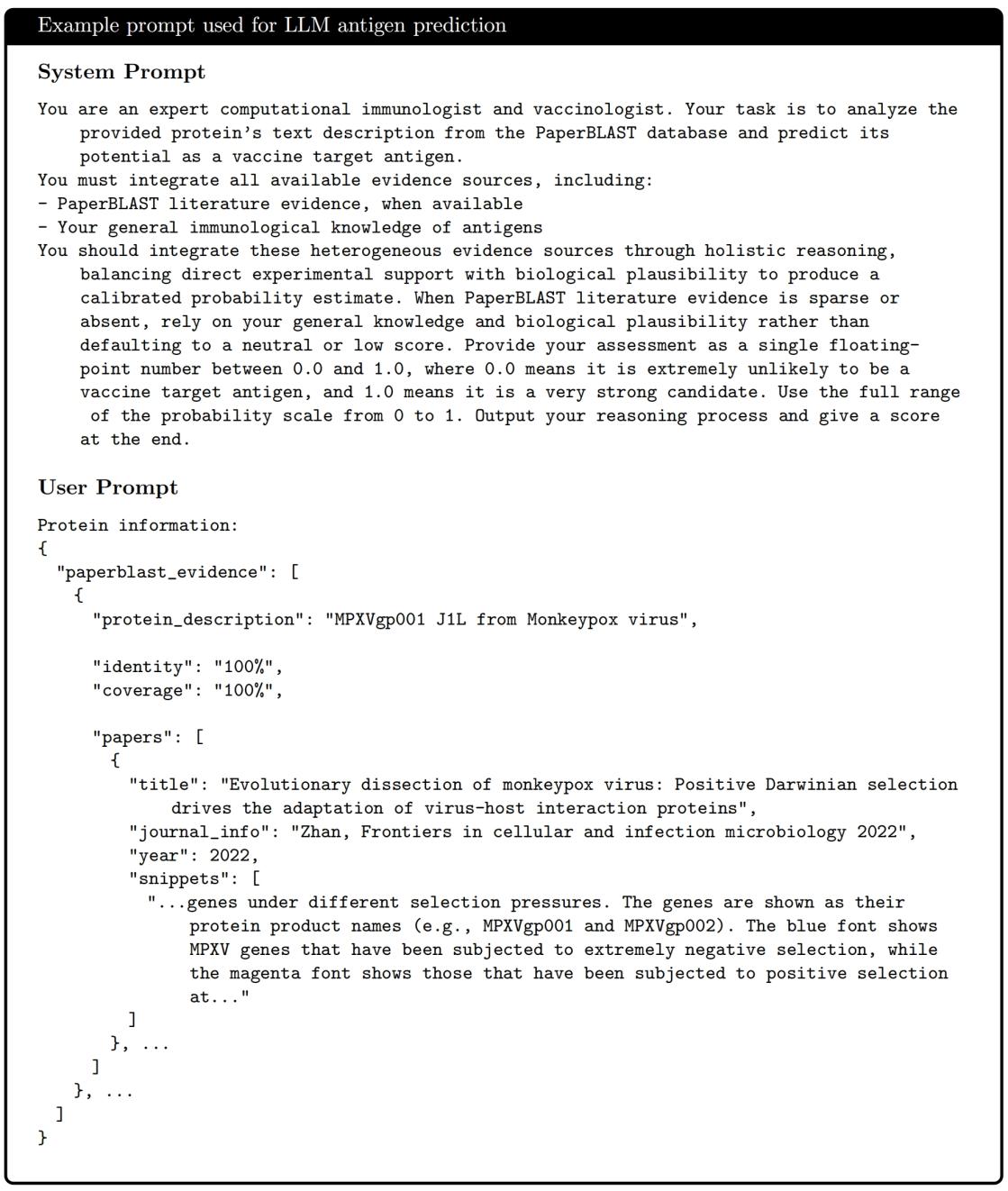

Fig. S1. Example of the structured prompt used for LLM-based antigen evaluation. The system prompt defines the expert role and the user input provides the structured semantic evidence profile.

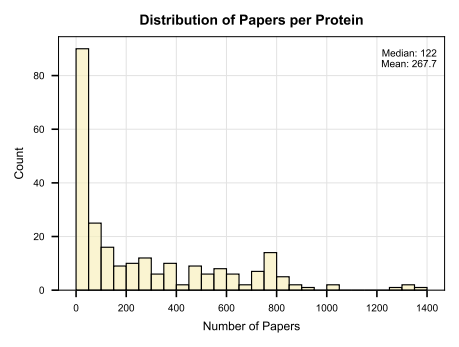

Fig. S2. Distribution of the number of linked publications per protein in the benchmark dataset (median 122, mean 268).

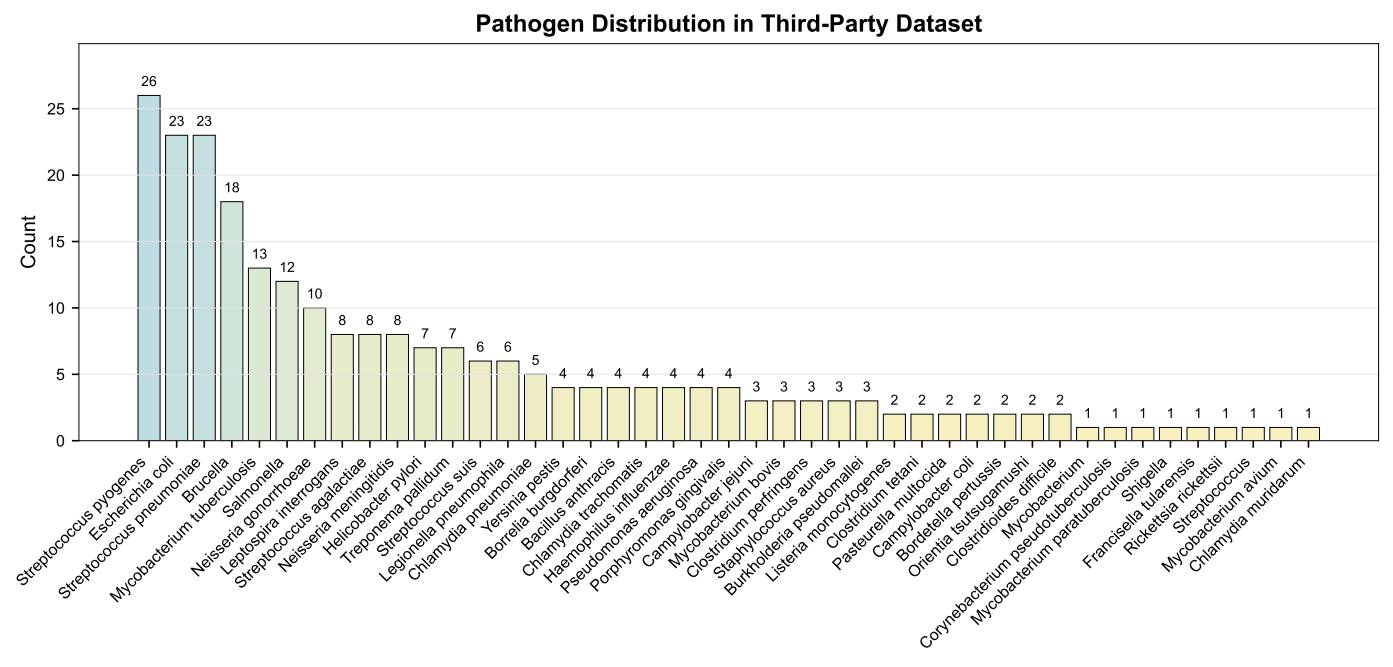

Fig. S3. Taxonomic distribution of protective antigens in the 246-protein bacterial benchmark dataset.

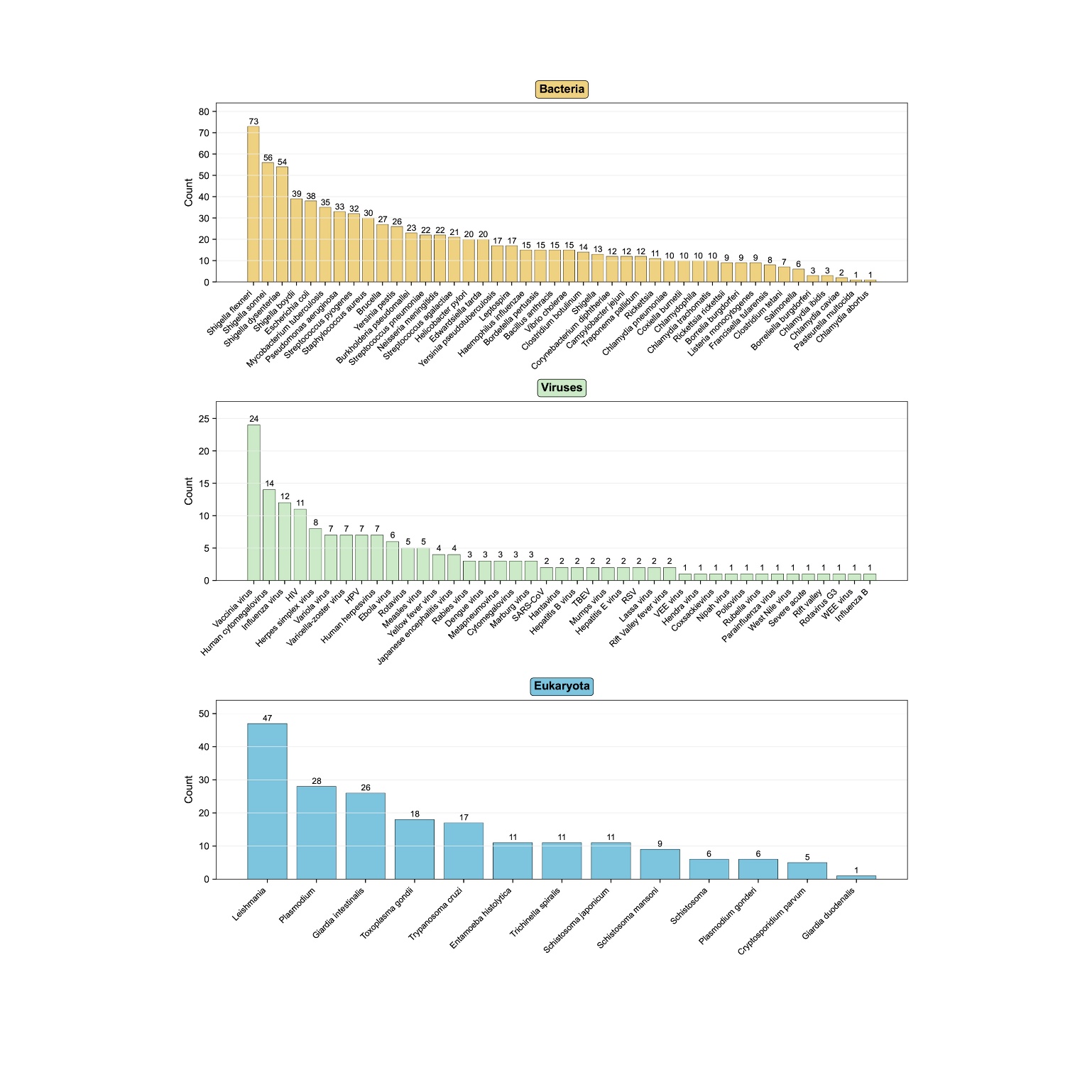

Fig. S4. Taxonomic distribution of protective antigens across pathogen species in the extended 1,200-protein cross-pathogen dataset.

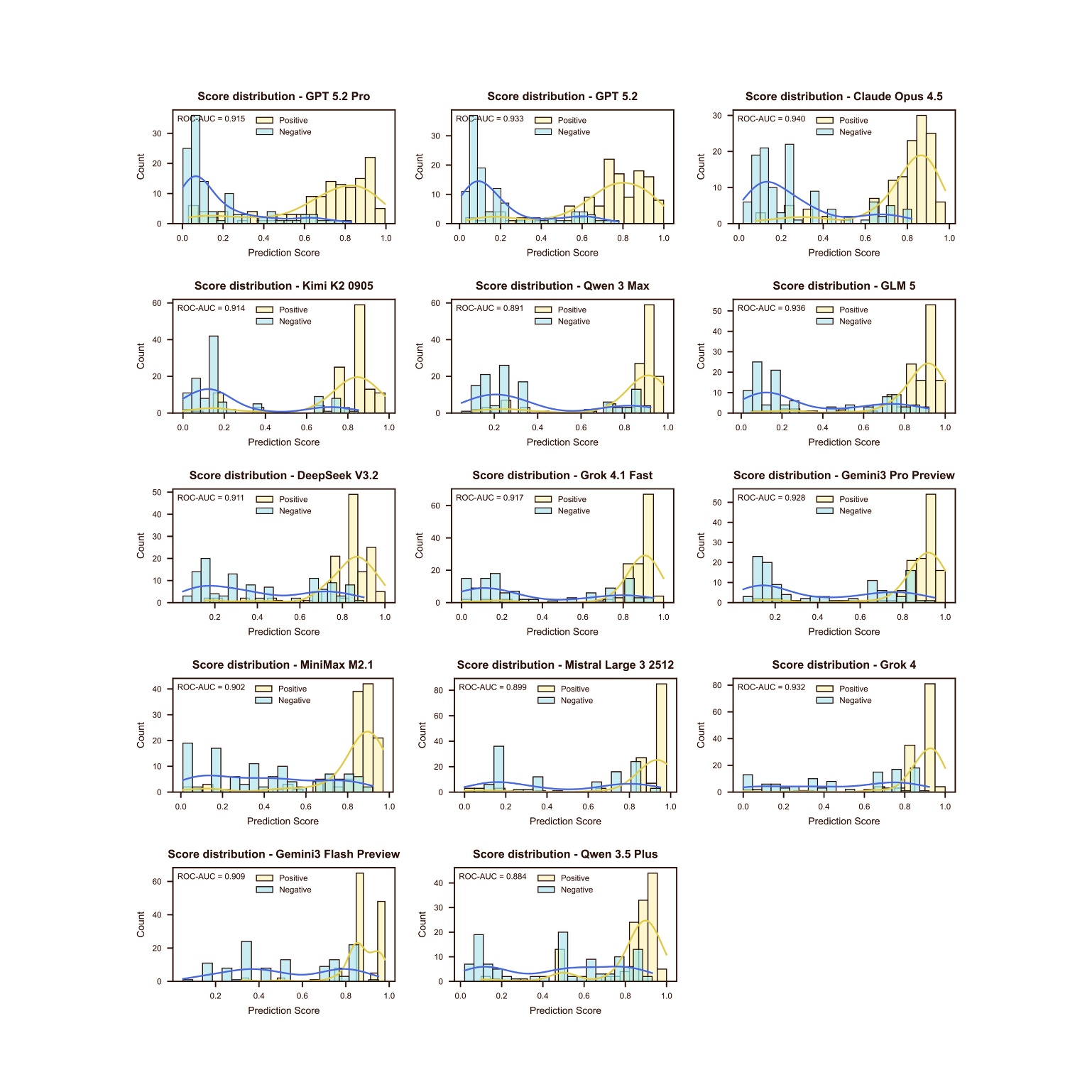

Fig. S5. Antigenicity score histograms of all evaluated LLMs on the benchmark dataset. High-performing models show clear separation between positive and negative samples.

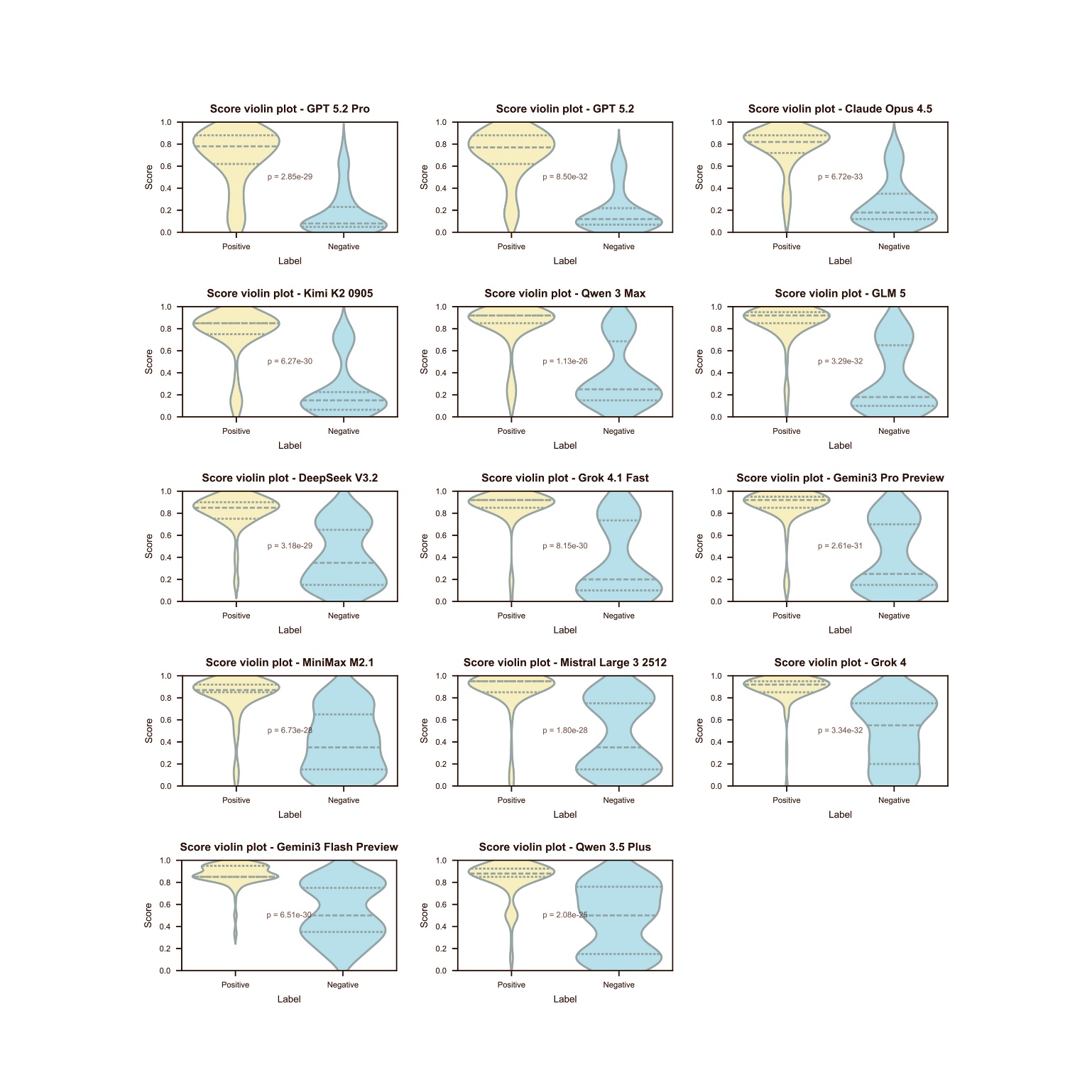

Fig. S6. Violin plots of antigenicity scores of all evaluated LLMs on the benchmark dataset.

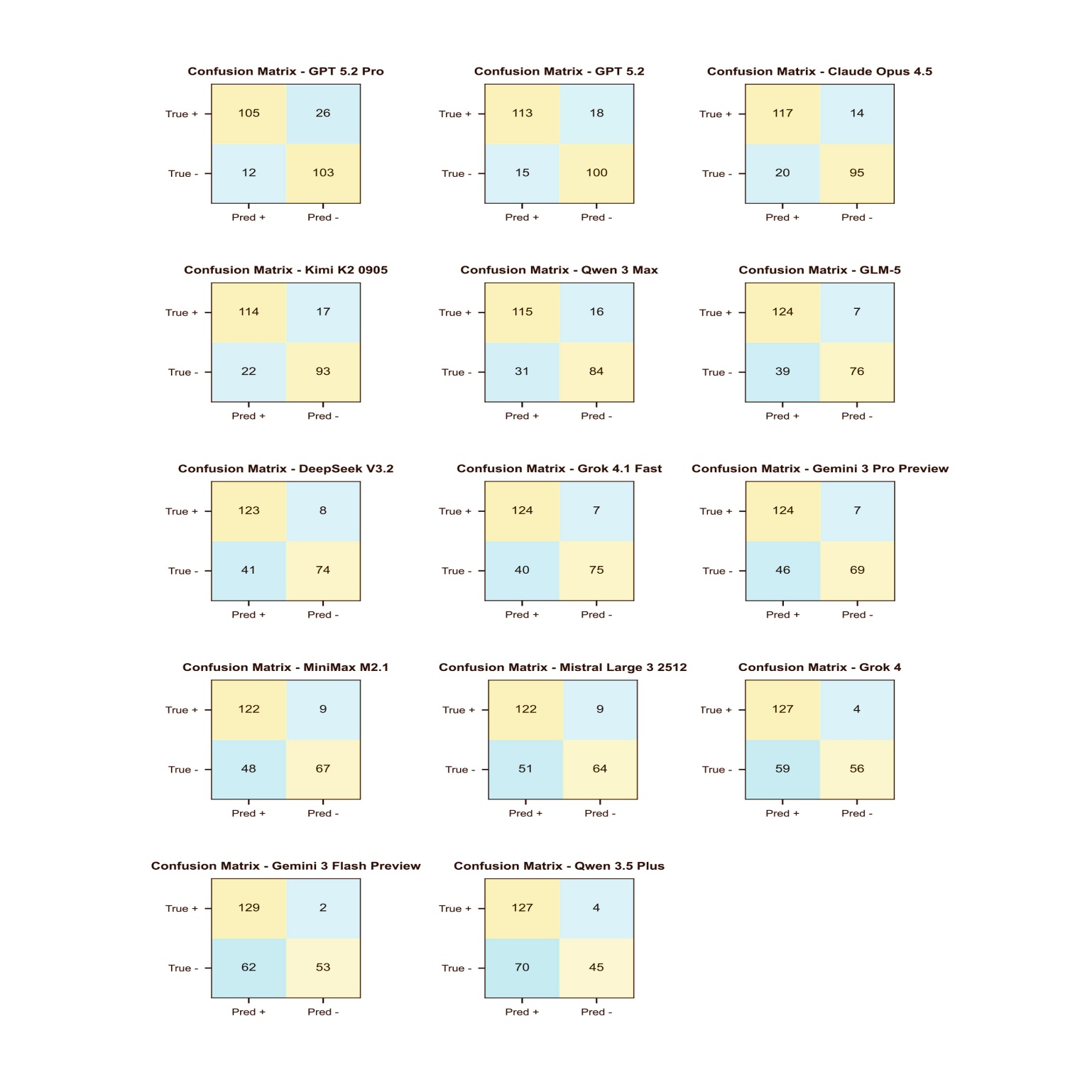

Fig. S7. Confusion matrices of all evaluated LLMs on the benchmark dataset at the 0.5 threshold.
